# Human gut microbiota networks disturbance by parasites in indigenous communities: Effect on bacteria genera related to depression incidence subnetworks

**DOI:** 10.1101/784470

**Authors:** Elvia Ramírez-Carrillo, Osiris Gaona, Javier Nieto, Andrés Sánchez-Quinto, Daniel Cerqueda-García, Luisa I. Falcon, Olga Rojas-Ramos, Isaac González-Santoyo

## Abstract

If you think you are in control of your behavior, think again. Evidence suggests that behavioral modifications, as development and persistence of depression, may be the consequence of a complex network of communication between macro (i.e. parasites) and micro-organisms capable of modifying the physiological axis of the host. Some parasites cause significant nutritional deficiencies for the host and impair the effectiveness of cognitive processes such as memory, teaching or non-verbal intelligence. Bacterial communities mediate the establishment of parasites and vice versa but this complexity approach remains little explored. We study the gut microbiota-parasite interactions using novel techniques of network analysis using data of individuals from two indigenous communities in the state of Guerrero, Mexico. Our results suggest that *A. Lumbricoides*, induce a gut microbiota perturbation affecting subnetworks of key species related to depression, consisting in the loss of network features such as path length, heterogeneity, number of nodes and neighbors; and especially by the loss of information emergence. Emergence is related with adaptability that has been linked to the concept of health as a critical balance between (adaptability) and self-organization (robustness). In this way, the loss of emergence means a depart from criticality and ultimately loss of health.

## Introduction

It is well documented that parasites can modulate several host’s behavioral patterns^1^, such as feeding, or reproductive behavior^2^. Some of these behavioral changes appear to directly benefit the parasite fitness, as is occurring with the trematode *Dicrocoelium dentriticum* and its host the ant *Formica fusca*, which modify its behavior even until death in order to accomplish the trematodes life cycle^3^. Nevertheless, other behavioral changes might be an incidental no-adaptive effect for the parasite, such as the host immunological response to rid the infection, which usually represent important physiological cost for the host^4^, or some parasite metabolic products that interfered with the communication pathways on the host’s central nervous system^5^. Hence, the physiological mechanisms involved in these behavioral changes being adaptive for the parasite or not, may include hormonal^6^, immunological^7^ and neurological components^8^. Nevertheless, these three levels of physiological signaling are not only regulated by parasites, but also by other microorganisms like bacterial microbiota, that coexist in the same host’s internal environment^9^.

In this sense, host’s behavioral changes might be viewed as the result of the complex communication network between macro and microorganisms that have the ability to modify these host’s physiological axis^9^. Moreover, the presence of certain bacterial communities should also impact the establishment of parasites and vice versa, parasites could be modifying the bacterial microbiota composition^9^. This bidirectional relation is plausible if both groups compete for similar host’s resources, such as a specific nutrient or an ecological niche, or because the activation of the host’s immune response due to the presence of parasites, disrupting different homeostatic relations established between bacterial microbiota and its host^10^.

In humans, *Ascaris lumbricoides* a soil-transmitted helminth (STH) affects more than a third of the world’s population, mainly in low-income populations in developing regions of Africa, Asia, and the Americas^11^. Its infection cause important nutritional deficits for the host^12^, and empirical evidence point out that it impairs the efficiency of cognitive processes, such as memory, learning or even non-verbal intelligence^13^. This helminth constantly interacts with other microorganisms, such as bacteria, archaea, yeasts, viruses, protozoa and even other helminths^14–17^.

In particular, the bacteria gut microbiota is the most diverse community of microorganisms with 1183 to 3180 genus reported so far^18^, and is undoubtedly essential to maintain the host health. For instance, to date at least 50 human pathologies have been associated with changes in the abundance and composition of gut microbiota^19^. While there are several well identified related pathologies include bowel disease, autoimmune diseases, metabolic syndromes, and neurological pathologies^19^; and second order effects by interactions, remains to be understood in depth.

These associations are due to the wide range of physiological functions they have. Human gut microbiota not only participates in most complex metabolic processes, such as fiber or starch catabolism, but also in the protection from pathogens^20–23^. It also intervenes in the proliferation and differentiation of our epithelium and it plays an important role in the development and modulation of the immune system^24^.

In recent years, it has also become evident that gut microbiota creates a two-way communication with the Central Nervous System (CNS). Such a relationship, known as the microbiota-gut-brain axis, has a deep impact on important brain mechanisms such as neuroinflammation, stress axis activation, neurotransmission and neurogenesis^25^. It has also been suggested that our gut microbiota composition may be affected by emotional variables such as stress or depression. For example, stress affects the intestinal epithelium and alters intestinal motility, its secretions and mucosa, thereby altering the bacterial habitat and affecting its composition and activity^26^. In addition, changes in the intestinal microbiota produce second-order effects and feedback that may affect motivation and other higher cognitive functions^25, 27^.

Recently relationship has been under quantitative study because advances in sequencing technology have made it possible to explore the role of the gut microbiota in a wide range of neurological and psychiatric disorders, including a larger scale analysis of self-reported conditions as performed by Valles-Colomer and his colleagues (2019)^28^ who investigate the gut microbiota compositional covariation with quality of life (QoL) indicators and general practitioner-reported depression in the Belgian Flemish Gut Flora Project (population cohort; n = 1,054).

In particular, they found that *Coprococcus* and *Dialister* genera were depleted in people important indicators of depression. Gut microbiota may be affected by several socio-economic and cultural contexts through diet changes especially when it implies different intake of animal protein, carbohydrates and processed food^29–31^; or medical practices like the over-ingestion of antibiotics^32^. Nevertheless, not only external context influence the gut microbiota ecosystems, but as was mentioned before the interaction with other microorganisms, as is occurring with helminth infections like *Ascaris lumbricoides*^17^. This might be a novel research field, a multi-ecosystemic perspective of microbiota-gut-brain axis, which remains little explored.

In this perspective, Chabé and coworkers (2017)^33^ discuss the possibility of a long coevolutionary history and tolerance for human parasites, since although some STH can cause severe illness^34, 35^; infections are often asymptomatic or in the case of many protozoa like Blastocystis spp. shows a high prevalence in microbiomes of healthy populations^36–38^.

Although different people, even in the same population, may present considerable microbial species variability, there has been recognized that gut microbiota exhibits some sort of ecological stability that translate into the fact that key species tends to remain present for long periods of time^23, 39^. This stability property of gut microbiota is considered key for host health and well-being, because it ensures that beneficial symbionts and their associated functions are maintained over time^40^.

In that sense healthy hosts should have gut microbiota in what has been called criticality, the balance between robustness and adaptability^41^. For instance, a healthy microbiota should have sufficient adaptation to respond to external variability, like changes on types of food available; but it also need to be robust in terms of key bacteria populations. The Criticality Hypothesis, states that systems in a dynamic regime shifting between order and disorder, attain the highest level of computational capabilities and achieve an optimal trade-off between robustness and adaptability^42^. In this framework robustness is associated with order and self-organization, meanwhile adaptability is related with disorder and information emergence, as we will discuss below. Empirical evidence has related human health to heart, and brain criticality^43–46^, and loss of criticality (mainly by loss of adaptability) with chronic diseases (such as obesity or diabetes) and elderly process^47^. In their work, Huitzil and co-workers (2018)^41^ claim that microbiome and genome networks are critical networks which means that their dynamical behavior is at the brink of a phase transition between order and chaos^48, 49^. This idea is supported by the facts that dynamical criticality confers the system properties such as evolvability (i.e., the coexistence of robustness and adaptability)^50, 51^, faster information storage, processing and transfer^52, 53^, and collective response to external stimuli without saturation^54^; and in fact there is solid evidence indicating that gene regulatory networks of real organisms are dynamically critical or close to criticality^55–58^.

This kind of multi-ecosystemic, complex system perspective present some serious challenges since microbiota contains many diverse species interacting with one another^40^, which makes the full system complex and challenging to understand. Network analysis has proven to be a valuable framework to understand large and complex interacting communities^59^. For instance, network analysis allows to study not only the whole ecosystem but also to focus on key bacteria for microbiota-gut-brain axis subnetworks (communities). In particular, Valles-Colomer and co-workers (2019)^28^ has reported specific gut bacteria genus related with wellbeing and depression. Therefore, in the present work at first, we explore the gut microbiota ecosystem considering whether the presence of the STH *A. Lumbricoides* predicts the gut microbiota network for adults and children (both female and male) in two poor indigenous non-industrialized communities with the highest index of STH infections in México. Secondly, we focus on how *A. Lumbricoides* infection is associated with subnetworks of bacteria communities strongly related with human depression symptomatology: *Coprococcus* and *Dialister* with a negatively relation.

## Results

We present analysis of The Graph Edit Distance (GED) analysis, which is a tool used for compare complete networks estructures. In Figure 1.A we show GED scores to compare Not parasitized (NP) vs Parasitized (P) populations divided by age group: Adults and Children. The scale goes from no difference (0) to totally different (1). Major differences are observed between Adults-NP Vs Adults-P that differ around 48%, followed by Children-NP Vs Adults-P that differ some 38% and finally Adults-P vs Children NP with a difference of 30%. The smallest differences (less affection from parasitosis) were found between Children-NP Vs Children-P. On the other hand, Adults showed greater affectation due to the presence of parasites. In order to compare the magnitude of differences, in Figure 1.B we show the GED scores paired, disaggregating data using age and gender. In all cases the magnitude of the difference is less than that caused by the presence of parasites.

**Figure 1.**
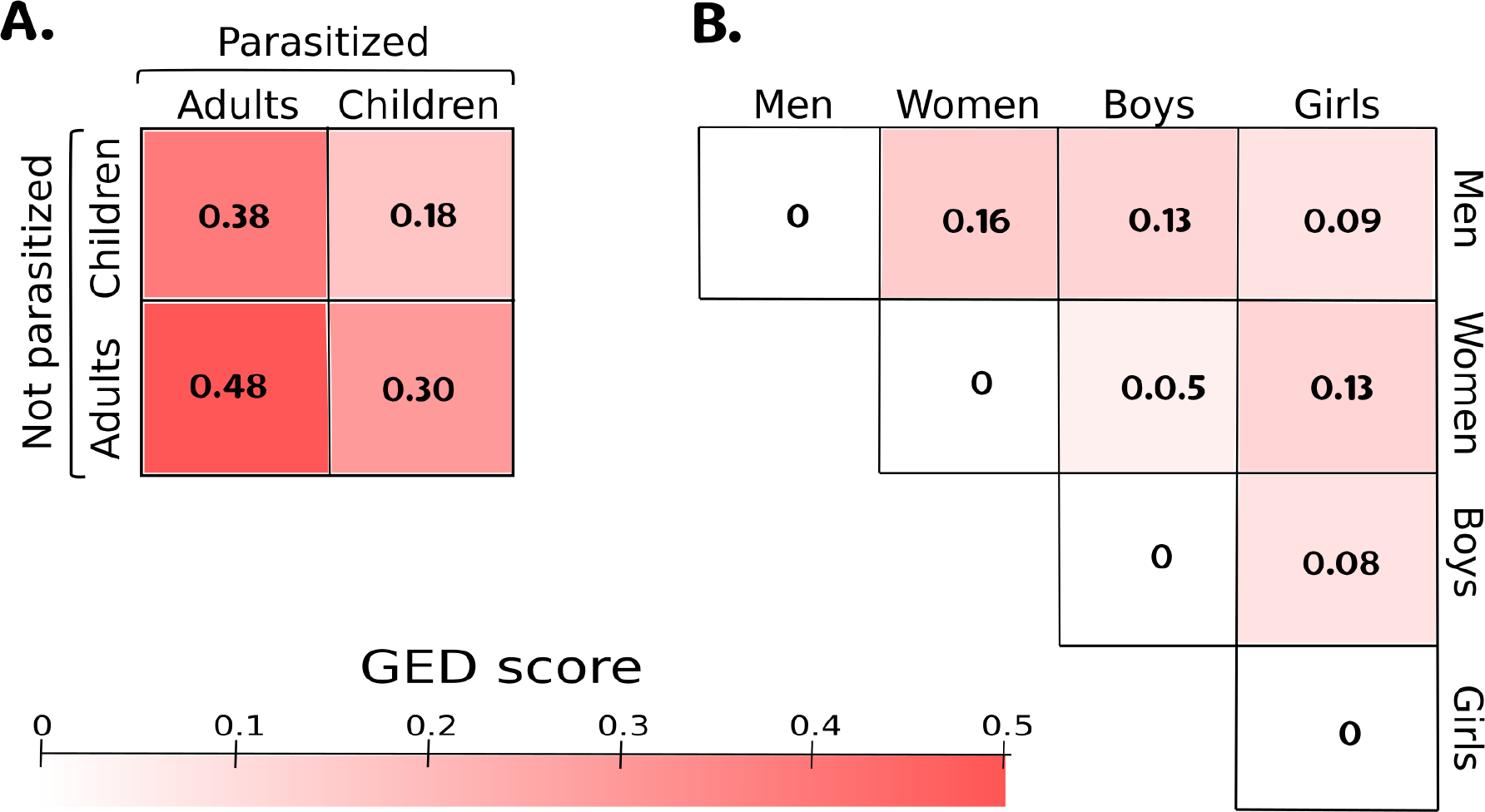
We show Graph Edit Distance (GED) comparing different pair of networks. The scale [0-1] goes from no difference to totally difference. (A) Show the comparisons between Not parasitized (NP) Vs Parasitized (P) populations divided by Adults and Children. (B) Differences between networks disaggregated by age and gender. The network of the microbiota in adults is more affected by the presence of parasites showing a 48% difference between treatments. Children are the least affected, differing only by 18%. In all cases the presence of parasites showed more differences when comparing the magnitudes by treatments than between ages or sexes.

Parasitosis also altered other standard network analysis measurements including Characteristic Path Length, Average number of neighbours, Number of nodes and Network heterogeneity (Fig.2). The parasitized adult and children networks show a decrease in all measures, which are related with different aspects of complexity which in turn may be defined^60^ as the product of emergence (adaptability) measured directed as Shannon Information (S) and self-organization (robustness) measured as its complement. So Complexity C= Emergence*Self-organization = S(1−S) = S−S2, is a quadratic form of information emergence meaning too little or too much emergence implies lower levels of complexity. Following the above, we calculated the networks Shannon Information, shown in Figure 3, which gives us information about the emergence of the systems. We found that Emergence is bigger in children than in adults (t=2.36, p=0.02) and as discussed this implies more adaptability. On the other hand, adults show a diminish in S, due to the presence of parasites (W=167, p=0.03).

**Figure 2.**
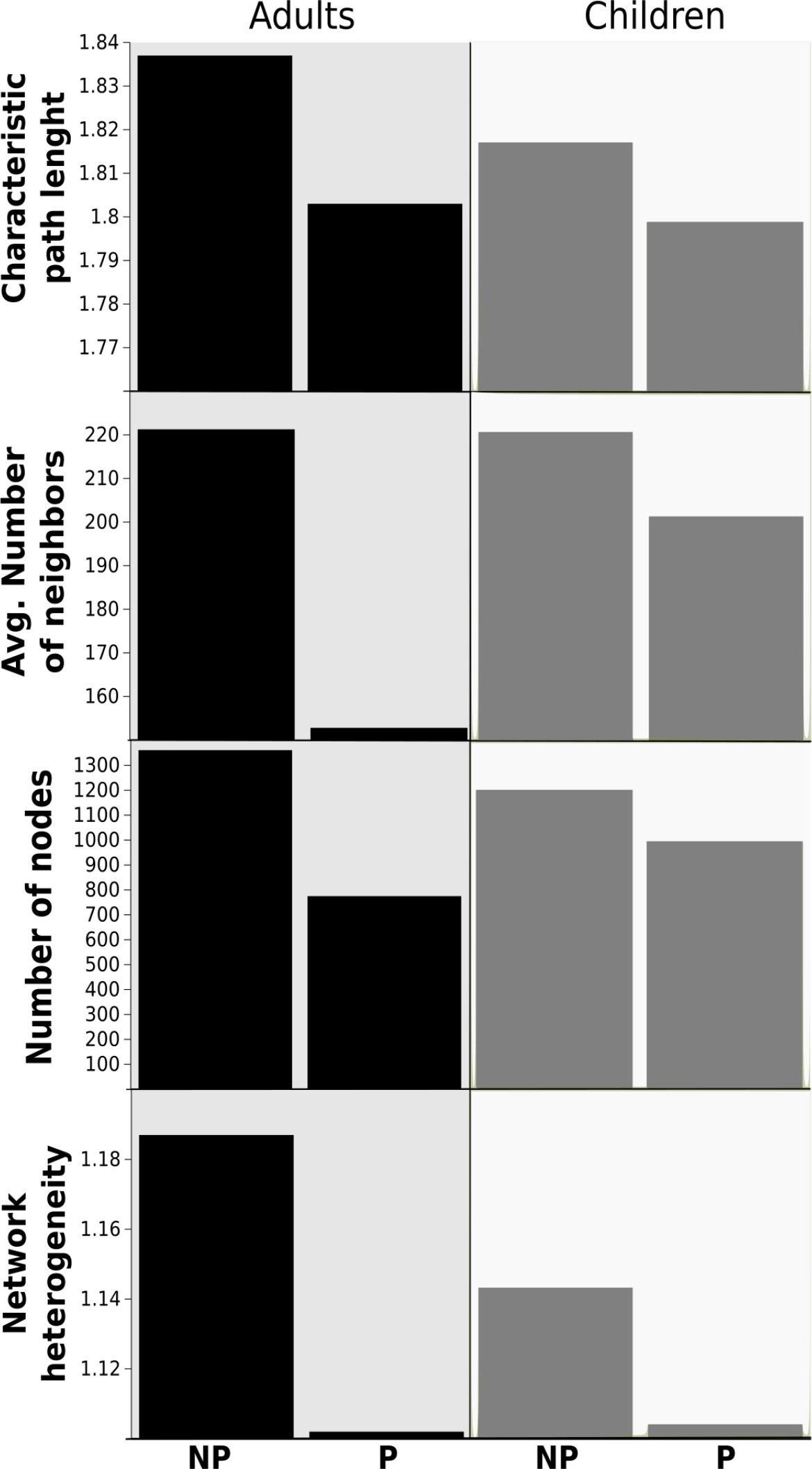
Standard network analysis measurements. In all the measurement network of individual parasitized both adults and children show a lower Characteristic Path Length, Average number of neighbours, Number of nodes and Network heterogeneity.

**Figure 3.**
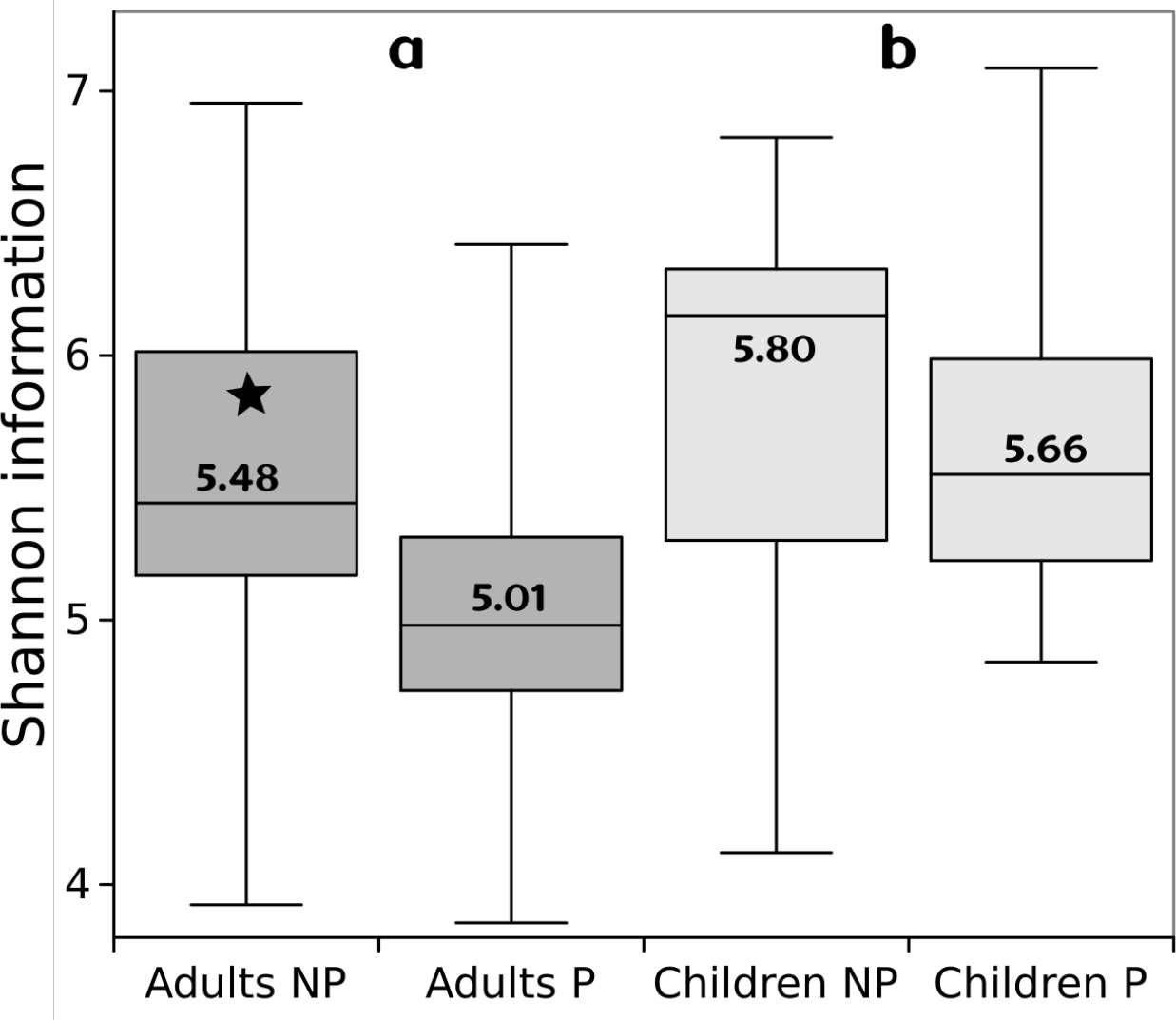
Shannon information as a measurement of information emergence. We found that Emergence is bigger in children than in adults (t=2.36, p=0.02) and as discussed this implies more adaptability. On the other hand, adults show a diminish in Shannon Information (S), due to the presence of parasites (W=167, p=0.03).

In turn, Bak and Paczuski (1995)^61^ has pointed out that Complexity arises from the tendency of large dynamic systems to become critical. Then as Complexity and Criticality are inherently connected^62^, a lost in Complexity translate in a depart form Criticality and most likely from healthy states too.

In order to understand in more depth the difference between networks we construct specific subnetworks for the most relevant species in terms of weight and connectivity, using the Maximal Clique Centrality (MCC) algorithm implemented in cytoHubba package^63^ who found MCC outperforms the other 11 methods. Figure 4 and 5 shows the 20 most important species according with MCC. Fig. 4 correspond to adults subnetwork and Fig.5 for children, in both cases with or without parasites. Inside the boxes are the species identifier number, listed below. Color encode phyla and star symbol which of them are present in both NP, P networks.

**Figure 4.**
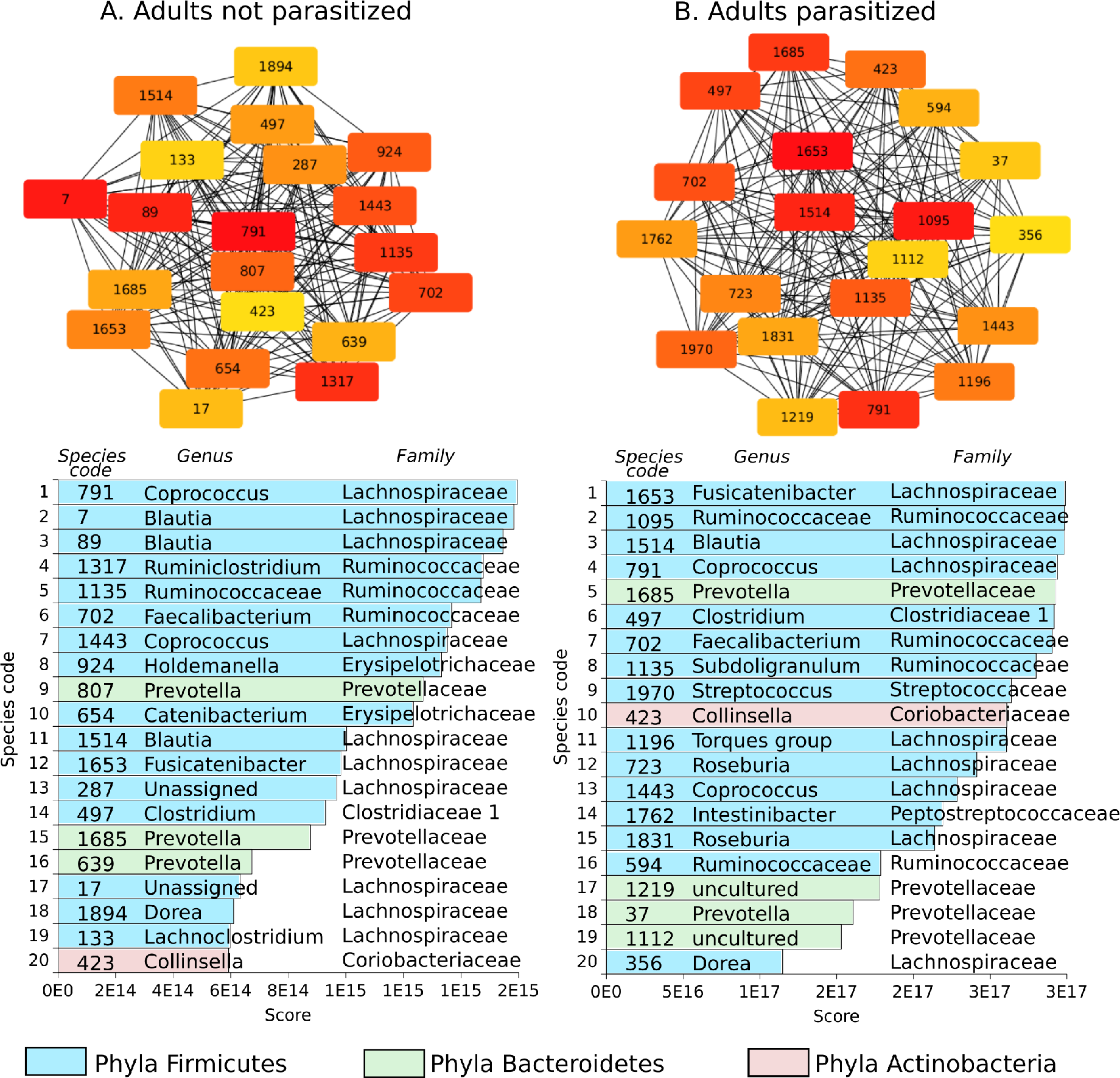
We show the 20 most important species for Adults-NP Vs Adults-P, calculated using the Maximal Clique Centrality (MCC) algorithm implemented in cytoHubba package^63^ who found MCC outperforms the other 11 methods. Inside the boxes are the species identifier number. The list shows each species score along side with genera, family an phyla. Color encode phyla and star symbol which of them are present in both NP, P networks.

**Figure 5.**
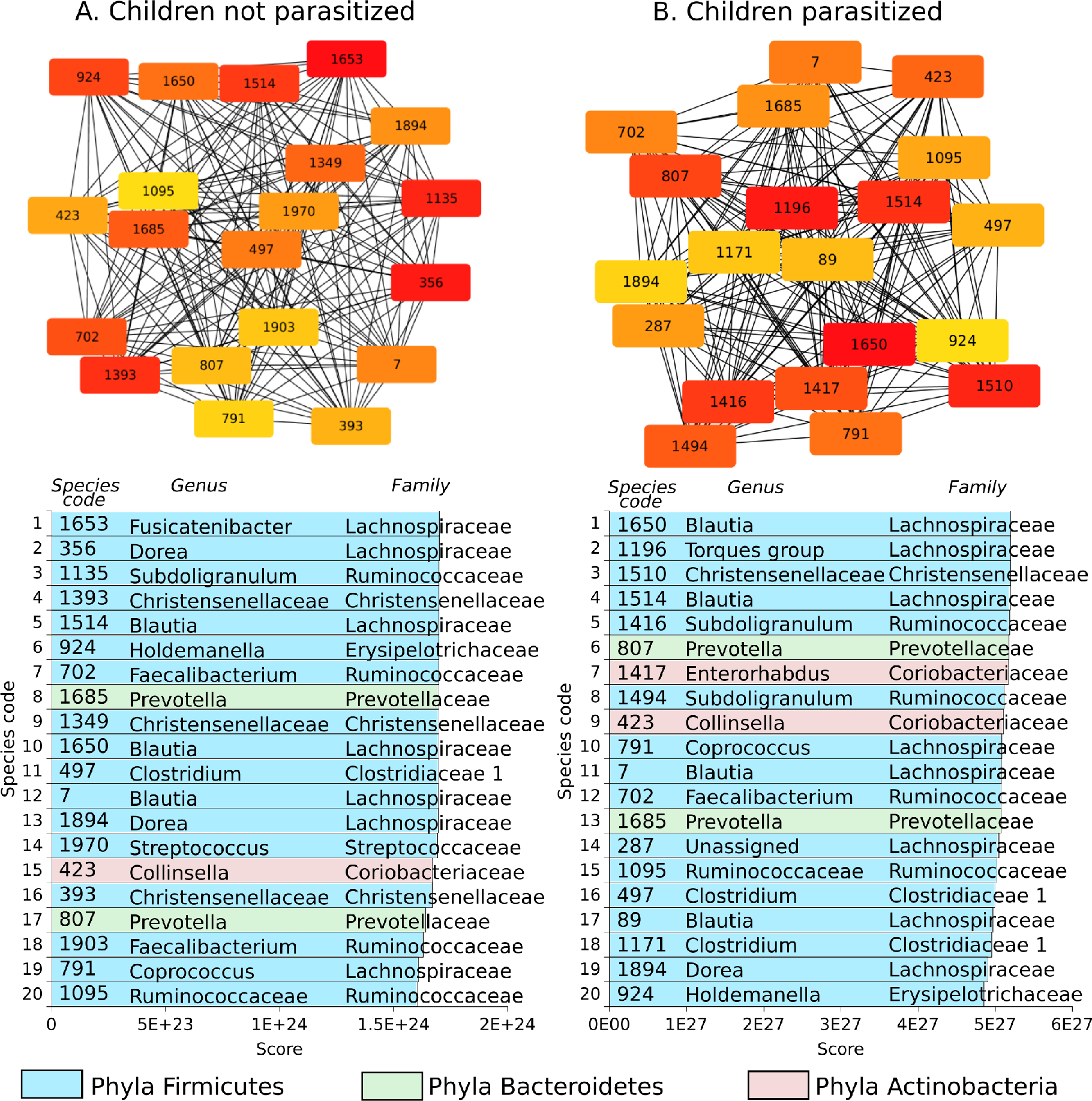
We show the 20 most important species for Children-NP Vs Children-P, calculated using the Maximal Clique Centrality (MCC) algorithm implemented in cytoHubba package^63^ who found MCC outperforms the other 11 methods. Inside the boxes are the species identifier number. The list shows each species score along side with genera, family an phyla. Color encode phyla and star symbol which of them are present in both NP, P networks.

We observe that although for complete networks are clear differences, it is not the case for more important MCC subnetworks, which makes sense since it has been acknowledged that gut microbiota has some kind of ecological stability which translates into the fact that important species tend to stay present for a long time^23, 39^

To see the effect of the parasites on the specific subnetworks of genus bacteria generates *Coprococcus* and *Dialister*. We compared the subnetworks of both species for adults and children parasitized and non-parasitized. A linear model was carried out to explain the variation of the wealth with respect to the presence of parasites, age and species (*R*^2^ = 0.78, p = 0.001) and a marginally significant negative relationship was found (t = −1.97, p = 0.05) between wealth and the presence of parasites. But beyond that the number of nodes (species) decreases with the presence of parasites, the results of the subnetworks shown in figures 6 and 7 allow us to observe how interactions with other species within the network are affected. For *Coprococcus*, children present more species richness than adults and in both cases, prasitosis reduces number of species, but it is interesting that in adults this link does not present links with other species, and in children the interactions are lost due to the presence of *A. Lumbricoides*. On the other hand, subnetworks of the *Dialister* genus are very interesting because adults are the ones that present a greater richness of species but with fewer interactions than in children. And in this case the presence of the parasite in adults collapses the network to only two species without interaction. In contrast, in children, although the number of species is also reduced, interactions increase and in fact interact with species from other new families in sub-networks such as the uncultured bacterium of Intestinibacter genus of Peptostreptococcaceae family.

**Figure 6.**
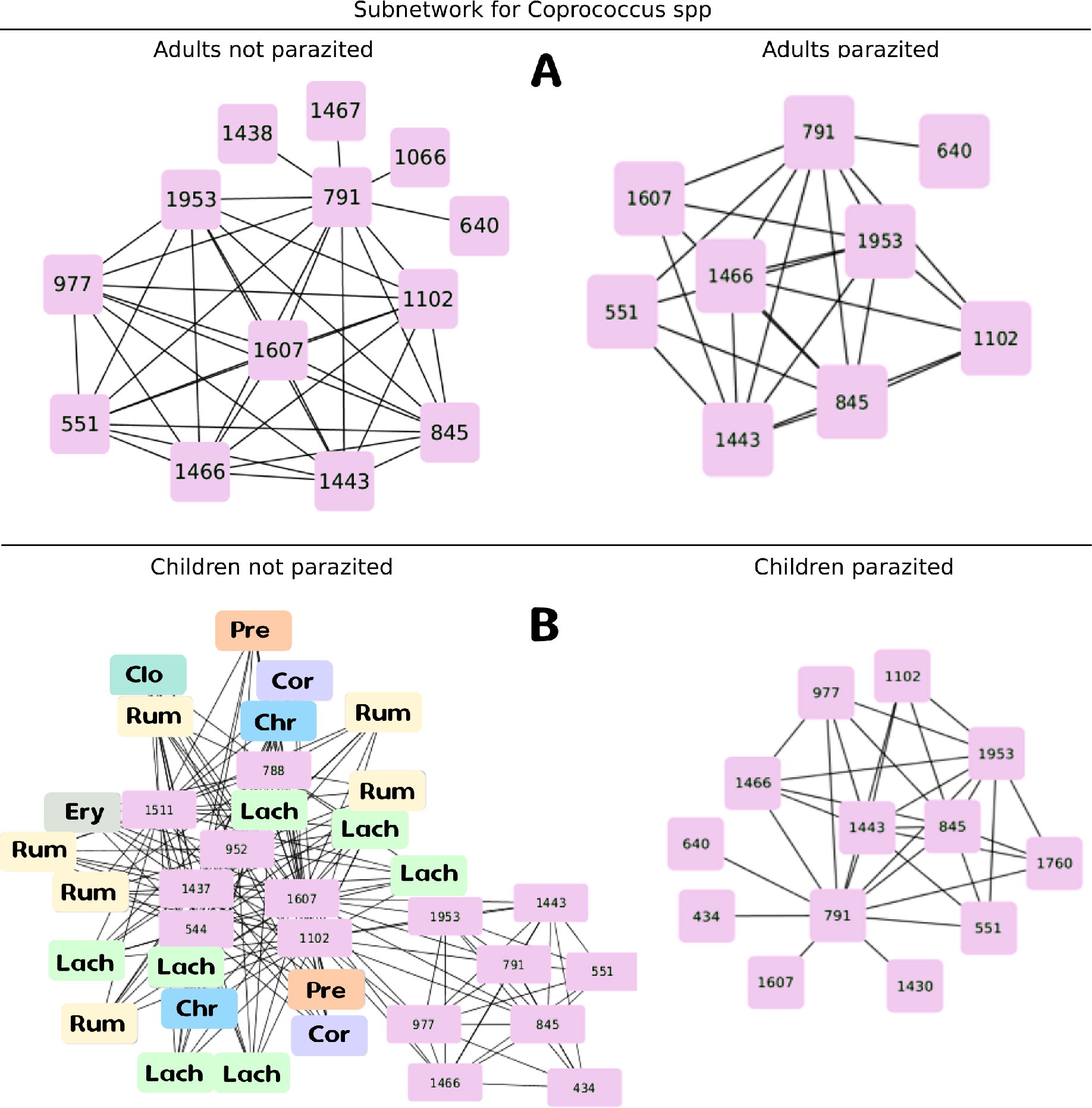
Subnetwork for bacteria species of *Coprococcus* genera, related to the incidence of depressive disorders as reported by (Valles-Colomer et al., 2019). Upper subfigure corresponds to adult population, in the left adults not parasitized (NP) and in the right adults parasitized (P). Lower subfigure corresponds to children populations, in the left children not parasitized (NP) and in the right children parasitized (P). In pink boxes are the species code for the diverse species of the genus *Coprococcus* founded in the subnetwork. In color boxes there are the species with which they interact in the network. the letter code indicates the families: Christensenellaceae (Chr), Clostridiaceae (Clo), Coriobacteriaceae (Cor), Erysipelatrichaceae (Ery), Lachnospiraceae (Lach), Peptostreptococcaceae (Pep), Prevotellaceae (Pre), Ruminococcaceae (Rum).

**Figure 7.**
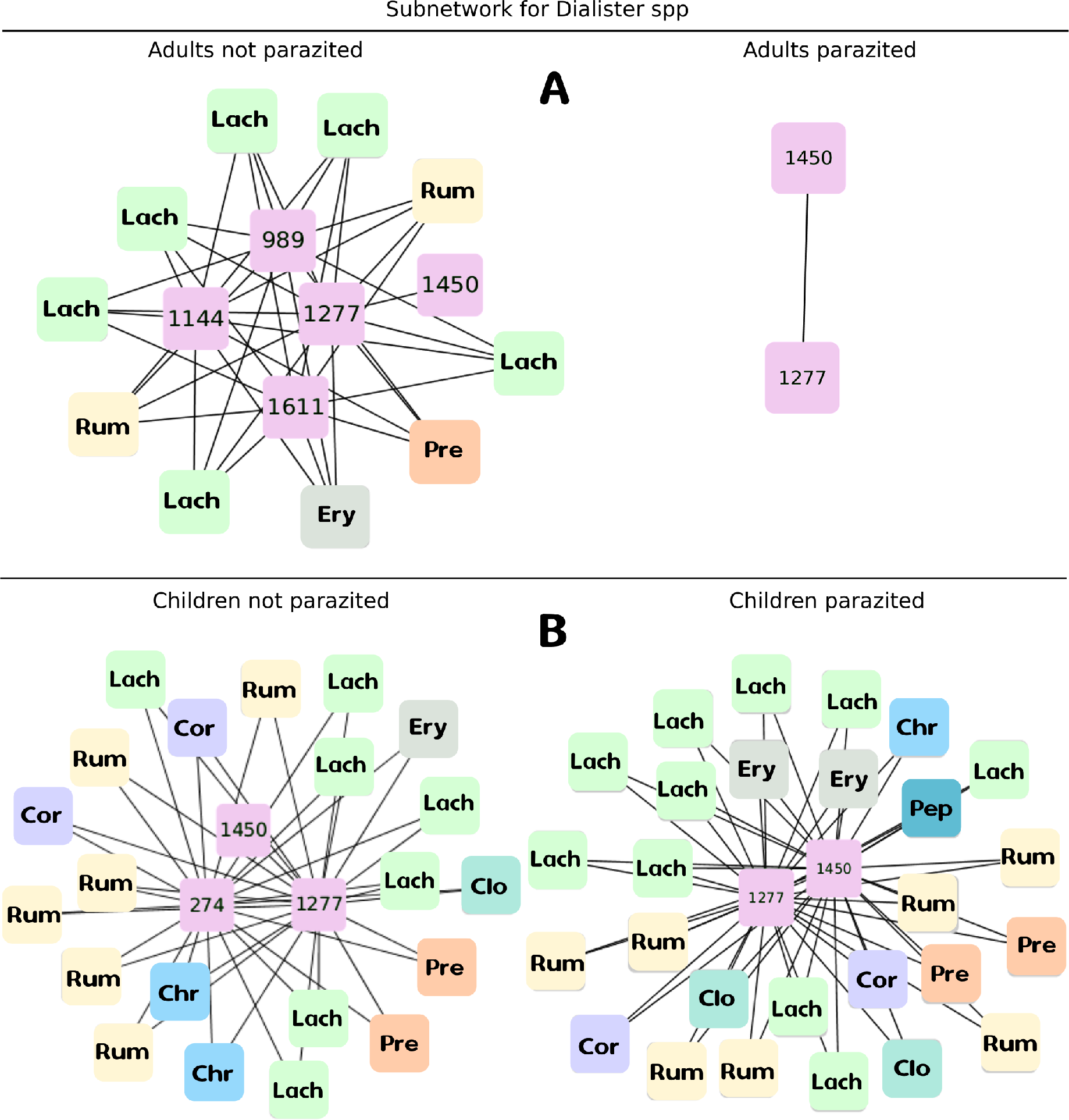
Subnetwork for bacteria species of *Dialister* genera, related to the incidence of depressive disorders as reported by (Valles-Colomer et al., 2019). Upper subfigure corresponds to adult population, in the left adults not parasitized (NP) and in the right adults parasitized (P). Lower subfigure corresponds to children populations, in the left children not parasitized (NP) and in the right children parasitized (P). In pink boxes are the species code for the diverse species of the genus *Coprococcus* founded in the subnetwork. In color boxes there are the species with which they interact in the network. The code of letters indicate the families with the same code of the previous figure

## Discussion

In here, we have studied from a complex systems perspective the effect of the STH *Ascaris lumbricoides* in the network properties of the host’s gut microbiota, focusing on particular effects of the disturbance on key bacteria genera that their absence is strongly related with depression: *Coprococcus* and *Dialister*.

The interaction of parasites with microbiota is an open hot field of study^28^ with complex interplays that reflects into both quality of life and depression. In the case of adults and children from the indigenous communities studied, we observed that the presence of *A. Lumbricoides* alters the structure of the gut microbiota networks, being more affected in adults who change by 48% in presence of this intestinal parasite. Moreover, adults have the least emergency (adaptability) and it decreases significantly when presenting *A. Lumbricoides*. On the other hand, the population of children that had initially greater diversity turns out to be more resistant and less affected to disturbances in the form of parasitism. An interesting question is whether our microbiota is losing criticality (via loss of adaptability) as we grow, as has been observed in the electrical activity of the heart^47^.

Interestingly, although the presence of *A. Lumbricoides* decreases the number of nodes (species) in complete networks, the analysis of the sub-networks of the 20 most relevant species (according to the MCC), presents relatively few changes. This may indicate that there is a certain “kernel” of species that are maintained despite the disturbances giving stability to the microbiota system^23, 39^.

The novelty of these analyses is that they allow us to analyze the interactions from a complex perspective, allowing us to see the whole system but also how different components are affected. In this sense, when analyzing the sub-networks for species related to depression, we observed that although in general species are lost, the interactions between them provide us with new insights. For example, Butyrate-producing species as *Coprococcus* sp. were consistently associated with a higher quality of life indicators^28^. Together with *Dialister*, *Coprococcus* genera were depleted in depression, even after correcting for confounding effects of antidepressants^28^.

We show that the presence of *A. Lumbricoides* impact in a particular way the subnetwork for these bacteria genera, first reducing the number of species that compose each genera in the net, and secondly reducing the interactions of these with other species. So, we can have different second-order effects by affecting interactions with other species. The way in which the interactions affect the species is complex and we still need to know a lot of particularity, but it could mean that the presence of parasites can promote relevant changes in networks of bacterial communities strongly related with the incidence of depression. These make sense, from a Criticality Hypothesis standpoint, with the lower values of network analysis measurements and Shannon Information for Parasites Adults and Children. These hypotheses pose that systems under criticality achieve an optimal trade-off between robustness and adaptability related with the highest level of computational capabilities. In particular, adaptability is related to emergence which in turn the highest level of computational capabilities and it can be understood as new global patterns, which are not present in the system’s components. More precisely, for continuous distributions, emergence interpretation is constrained to the average uncertainty a process produces under a specific set of the distribution parameters (e.g., the SD value for a Gaussian distribution), and so it can be measured using Shannon information as proposed by Santamiaría-Bonfil and co-workers (2017)^64^. Inhere we follow this interpretation of emergence and take it as Shannon Information.

Recent results show convincing evidence that criticality may be a key feature of a healthy state^43–46, 65^. Our results show that due to the presence of parasites, there is a depart from criticality via a diminish of emergence (adaptability) and then a loss of health. Nevertheless, the net effect of parasites interacting with microbiota maybe not as straightforward as some recent studies suggest^40^, due to the fact that we have been co-evolving with them and some types and intensity of parasitism might impose some sort of stressor for the microbiota, which may produce a hormesis effect contributing to healthier states. This second order effect relates to Taleb’s ideas^66^ about antifragility in medicine. Antifragility is a property that enhances the capability of a system to respond to external stressors in a nonlinear convex manner in the payoff space. Antifragile systems take advantage of volatility and stressors, so in the absence, this property could be lost.

## Methods

### Study site

In Mexico there are at least 56 independent indigenous peoples, whose lifestyle practices differ in varying degrees from the typical “westernized” lifestyle. Among these groups, the Me’Phaa people, from a region known as the “Montaña Alta” of the state of Guerero, is one of the groups whose lifestyle differs most strongly from the westernized lifestyle typical of more urbanized areas. In these communities there is almost no access to allopathic medications, and there is no health service, plumbing, or system of water purification. Water for washing and drinking is obtained from small wells. Therefore, these communities represent the lowest income in the country, the highest index of child and adult morbidity and mortality by intestinal infection (children’s age from 0 to 8 years old, which is the highest vulnerability and death risk age^67^), and the lowest access to health services. These conditions were determined by last 10 years of statistical information obtained from National information system of access to health^68^. Most Me’Phaa speak only their native language, and the closest large town (the main municipal town) is two hours away by dirty-road. Our data were collected from two of these indigenous communities; Plan de Gatica and El Naranjo. Distance between these two communities is about 30 km^69^, and their socio-ecomonic a cultural patterns are the same between them^69^. Although allopathic medication is practically absent in these communities^68^, we selected only participants that have not taken any medications during the last two years prior to study, such as antibiotics or anthelmintic treatment. Samples were taken from 63 individuals in total-35 from Plan de Gatica and 28 from El Narajo. Children were aged 5 to 10 years old, and adults were between 18 and 45. We sampled 29 children in total: 16 (7 from Gatica and 9 from Naranjo) and 13 (8 from Gatica and 5 from Naranjo), whose average age was 7.6 +/− 1.8 years. Among adults, we sampled 34 total. 18 were women (10 form Gatica and 8 from Naranjo) and 16 were men (10 from Gatica and 6 from Naranjo). The average age was 30.48 +/− 7.79 years.

### Extraction of DNA from feces

In order to characterize the abundance and composition of the participants’ intestinal microbiota, we used a High throughput sequencing method using 16 S ribosomal amplicons and mass sequencing, using an illumine platform. Therefore, two grams of fecal samples from each participant were received in sterile containers, then transferred to 1.5 milliliter Eppendorf tubes using sterile technique, and transported in liquid nitrogen at negative 80 degrees Celsius until DNA extraction. Metagenomic fecal DNA was extracted using the DNeasy Blood & Tissue kit (Qiagen, Valencia, CA) according to the manufacturer’s instructions. Briefly, feces collected into 1.5 ml sterile tubes were diluted with 180 *μ*l of ATL extraction buffer with 20 *μ*l proteinase K (10 mg ml-1). Tubes were mixed thoroughly by vortexing and were incubated at 56°C at 1500 rpm for 50 min. 200 *μ*l of AL Buffer with 200 *μ*l ethanol (96-100%) were added and mixed thoroughly by vortexing. The mixture was transferred into the DNeasy Mini spin column, washed with Buffer AW1 and then with AW2. The DNA was eluted with 200 *μ*l of AE Buffer and precipitated with absolute ethanol, 0.1 volume 3 M sodium acetate and 2 *μ*l glycoblue. DNA was resuspended in 30 *μ*l of molecular grade water and stored at −20°C until PCR amplification.

### 16S rRNA gene amplification and sequencing

DNA samples were PCR-amplified using the hypervariable V4 region of the 16S rRNA gene with universal bacteria/archaeal primers 515F/806R following the procedures reported by Caporaso et al. (2010)^70^ and Carrillo et al. (2015)^71^. PCR reactions (25 *μ*l) contained 2-6 ng of total DNA, 2.5 *μ*l Takara ExTaq PCR buffer 10X, 2 *μ*l Takara dNTP mix (2.5 mM), 0.7 *μ*l bovine serum albumin (BSA, 20 mg ml-1), 1 *μ*l primers (10 *μ*M), 0.125 *μ*l Takara Ex Taq DNA Polymerase (5 U *μ*l-1) (TaKaRa, Shiga, Japan) and nuclease-free water. Samples were amplified in triplicate using a PCR protocol including an initial denaturation step at 95°C (3 min), followed by 35 cycles of 95°C (30 s), 52°C (40 s) and 72°C (90 s), followed by a final extension (72°C, 12 min). Triplicates were then pooled and purified using the SPRI magnetic bead, AgencourtAMPure XP PCR purification system (Beckman Coulter, Brea, CA, USA). The purified 16S rRNA fragments (∼20 ng per sample) were sequenced on an IlluminaMiSeq platform (Yale Center for Genome Analysis, CT, USA), generating ∼250 bp paired-end reads. The sequence data are available from the NCBI Bioproject number PRJNA508738.

### Analysis of the sequence data

The paired-end 2×250 reads were processed in QIIME2^72^. The reads were denoised with the DADA2 plugin to resolve the amplicon sequence variants (ASVs). Both forward- and reverse-reads were truncated at 200 pb, and chimeric sequences were removed using the “consensus” method. Representative ASV sequences were taxonomically assigned using the “classify-consensus-vsearch pluggin”, using the SILVA 132 database as a reference. An alignment was performed with the MAFFT algorithm. After masking positional conservations and gap filtering, a phylogeny was built with the FastTree algorithm. The abundance table and phylogeny were exported to the R environment to perform the statistical analysis with the phyloseq, vegan and ggplot2 packages. Plastidic ASVs were filtered out of the samples (for subsequent separate analysis), then the samples were rarefied to a minimum sequencing effort of 10 000. Counts of plastidic ASVs (filtered before rarefaction) were normalized with the cumulative sum scaling (CSS) method with the metagenomeSeq package^73^. The total diversity (alpha diversity) of the ASVs was calculated using Shannon’s Diversity Index and Faith’s Phylogenetic Diversity.

### Determination of *A. Lumbricoides* presence

The presence of *A. Lumbricoides* was done in the same participant’s fecal samples used to determine the composition and abundance of its microbiota. The identification of this nematode was done through light field microscopy following the protocol of “Mini-FLOTAC”, standardized by Cringoli et al. (2010)^74^. This “Mini-FLOTAC” protocol is a novel and more sensitive quantitative method for the identification of STH compared to other techniques, such as Kato-Katz^75^.

### Network and subnetworks analyses

From the dataset of ASVs (i.e. bacteria species) relative abundances in fecal samples, we used the Cooccur package (https://cran.r-project.org/web/packages/cooccur/cooccur.pdf) in R^76^ to construct a co-occurrence matrix and use it as weight in networks for: Adults and Children with and without parasites.

Then we used CytoScape an open source software platform for visualizing and analyze complex networks^77^. First we used CytoHubb a CytoScape’s app which explore important nodes/hubs and fragile motifs in an interactome network by several topological algorithms including Degree, Edge Percolated Component (EPC), Maximum Neighborhood Component (MNC) among others to rank nodes (species). In particular we retain the 20 most important species using Maximal Clique Centrality (MCC) which has been reported to be the best option for this^63^.

Once ranked we construct sub networks and compare them in terms of species composition. In a recent work in Nature^28^ the authors report specific gut bacteria related with depression, which we look for in our data and construct subnetworks for each one. Beyond a composition analysis, we used an implementation of Graph Edit Distance (GED) a measure of similarity (or dissimilarity) between two graphs in the CytoGEDEVO app^78^. The GEDEVO^79^, method uses an evolutionary algorithm to iteratively optimize a number of random initial alignments. Candidate alignments are modified using mutation and crossover operations to yield better scoring variants after each iteration. The algorithm terminates after convergence is detected. In short, GEDEVO is a method for global topological graph alignment that minimizes graph edit distance (GED).

We used CytoGEDEVO to calculate the pared distance between fathers, mothers, daughters and sons. And then between children (male and female) with and without parasites; parents (male and female) with and without parasites.

### Ethical compliance

All study procedures are compliant with all relevant ethical regulations. All procedures for testing and recruitment were approved by the National Autonomous University of Mexico Committee on Research Ethics (FPSI/CE/01/2016), and run in accordance with the ethical principles and guidelines of the Official Mexican Law (NOM-012-SSA3-2012). Written informed consent was obtained from all participants. For all under 18 years, specific written informed consent was obtained from their parents or legal guardians according with ethical guidance and Committee.

## Data availability

The datasets generated during and/or analyzed during the current study are available from the corresponding author on reasonable request.

## Additional Information

We declare that the authors have no competing interests as defined by Nature Research, or other interests that might be perceived to influence the results and/or discussion reported in this paper.

## Acknowledgements

This project was funded by UNAM-PAPIIT [grant numbers IA209416, IA207019] and CONACYT Ciencia Básica [grant number 241744]. ER-C thanks PROGRAMA DE BECAS POSDOCTORALES EN LA UNAM (DGAPA/UNAM Posdoctoral fellowship program)

We are grateful to Xuajin Me’Phaa, Margarita Muciño, Julio Gatica, and Diego Hernandez-Muciño for their help in the liaison with the Me’Phaa community, and for their help in data collection logistics. We also are grateful to Santiago Martinez-Correa and Aida Elizondo García for their help in DNA amplification and parasite determination respectively.

## Author contributions statement

ER-C, JN, and IG-S, conceived and designed this study. AS-Q, OG, DC-G, LF, OR-R and IG-S collected data, including DNA extraction and amplification. ER-C, AS-Q, OG, DC-G, and IG-S analysed all data. All authors contributed in writing the first draft. All authors approved the final version of the manuscript and gave approval for publication.

## Notes

#### Summary of Updates

Non since there was no decision letter

## References

1. Adamo, S. A. Modulating the modulators: parasites, neuromodulators and host behavioral change. Brain, behavior evolution 60, 370–377 (2002).

2. González-Tokman, D., Córdoba-Aguilar, A., González-Santoyo, I. & Lanz-Mendoza, H. Infection effects on feeding and territorial behaviour in a predatory insect in the wild. Animal Behav. 81, 1185–1194 (2011).

3. Libersat, F., Emanuel, S. & Kaiser, M. Mind control: how parasites manipulate cognitive functions in their insect hosts. Front. psychology 9, 572 (2018).

4. Adamo, S. The specificity of behavioral fever in the cricket acheta domesticus. The J. parasitology 529–533 (1998).

5. Adalid-Peralta, L., Sáenz, B., Fragoso, G. & Cárdenas, G. Understanding host–parasite relationship: the immune central nervous system microenvironment and its effect on brain infections. Parasitology 145, 988–999 (2018).

6. Romano, M. C., Jiménez, P., Miranda, C. & Valdez, R. A. Parasites and steroid hormones: corticosteroid and sex steroid synthesis, their role in the parasite physiology and development. Front. neuroscience 9, 224 (2015).

7. Shepherd, C. et al. Identifying the immunomodulatory components of helminths. Parasite immunology 37, 293–303 (2015).

8. Johnson, T. P. & Nath, A. Neurological syndromes driven by postinfectious processes or unrecognized persistent infections. Curr. opinion neurology 31, 318–324 (2018).

9. Leung, J. M., Graham, A. L. & Knowles, S. C. Parasite-microbiota interactions with the vertebrate gut: synthesis through an ecological lens. Front. Microbiol. 9(2018).

10. Cortés, A., Toledo, R. & Cantacessi, C. Classic models for new perspectives: delving into helminth–microbiota–immune system interactions. Trends parasitology 34, 640–654 (2018).

11. Torgerson, P. R. et al. World health organization estimates of the global and regional disease burden of 11 foodborne parasitic diseases, 2010: a data synthesis. PLoS medicine 12, e1001920 (2015).

12. Hall, A., Hewitt, G., Tuffrey, V. & De Silva, N. A review and meta-analysis of the impact of intestinal worms on child growth and nutrition. Matern. & child nutrition 4, 118–236 (2008).

13. Guernier, V. et al. Gut microbiota disturbance during helminth infection: can it affect cognition and behaviour of children? BMC infectious diseases 17, 58 (2017).

14. Eckburg, P. B. et al. Diversity of the human intestinal microbial flora. science 308, 1635–1638 (2005).

15. Gaci, N., Borrel, G., Tottey, W., O’Toole, P. W. & Brugère, J.-F. Archaea and the human gut: new beginning of an old story. World journal gastroenterology: WJG 20, 16062 (2014).

16. Scarpellini, E. et al. The human gut microbiota and virome: Potential therapeutic implications. Dig. Liver Dis. 47, 1007–1012 (2015).

17. Williamson, L. L. et al. Got worms? perinatal exposure to helminths prevents persistent immune sensitization and cognitive dysfunction induced by early-life infection. Brain, behavior, immunity 51, 14–28 (2016).

18. Falony, G. et al. Population-level analysis of gut microbiome variation. Science 352, 560–564 (2016).

19. Moya, A. & Ferrer, M. Functional redundancy-induced stability of gut microbiota subjected to disturbance. Trends microbiology 24, 402–413 (2016).

20. Bäckhed, F., Ley, R. E., Sonnenburg, J. L., Peterson, D. A. & Gordon, J. I. Host-bacterial mutualism in the human intestine. science 307, 1915–1920 (2005).

21. Mazmanian, S. K., Round, J. L. & Kasper, D. L. A microbial symbiosis factor prevents intestinal inflammatory disease. Nature 453, 620 (2008).

22. Lee, Y. K. & Mazmanian, S. K. Has the microbiota played a critical role in the evolution of the adaptive immune system? Science 330, 1768–1773 (2010).

23. Dethlefsen, L. & Relman, D. A. Incomplete recovery and individualized responses of the human distal gut microbiota to repeated antibiotic perturbation. Proc. Natl. Acad. Sci. 108, 4554–4561 (2011).

24. Rojo, D. et al. Exploring the human microbiome from multiple perspectives: factors altering its composition and function. FEMS microbiology reviews 41, 453–478 (2017).

25. Sherwin, E., Sandhu, K. V., Dinan, T. G. & Cryan, J. F. May the force be with you: the light and dark sides of the microbiota–gut–brain axis in neuropsychiatry. CNS drugs 30, 1019–1041 (2016).

26. Collins, S. M. & Bercik, P. The relationship between intestinal microbiota and the central nervous system in normal gastrointestinal function and disease. Gastroenterology 136, 2003–2014 (2009).

27. Mayer, E. A., Savidge, T. & Shulman, R. J. Brain–gut microbiome interactions and functional bowel disorders. Gastroenterology 146, 1500–1512 (2014).

28. Valles-Colomer, M. et al. The neuroactive potential of the human gut microbiota in quality of life and depression. Nat. microbiology 1 (2019).

29. Wu, G. D. et al. Linking long-term dietary patterns with gut microbial enterotypes. Science 334, 105–108 (2011).

30. David, L. A. et al. Diet rapidly and reproducibly alters the human gut microbiome. Nature 505, 559 (2014).

31. Xu, Z. & Knight, R. Dietary effects on human gut microbiome diversity. Br. J. Nutr. 113, S1–S5 (2015).

32. Francino, M. Antibiotics and the human gut microbiome: dysbioses and accumulation of resistances. Front. microbiology 6, 1543 (2016).

33. Chabé, M., Lokmer, A. & Ségurel, L. Gut protozoa: friends or foes of the human gut microbiota? Trends Parasitol. 33, 925–934 (2017).

34. Cwiklinski, K., O’neill, S., Donnelly, S. & Dalton, J. A prospective view of animal and human fasciolosis. Parasite immunology 38, 558–568 (2016).

35. Mutapi, F. The gut microbiome in the helminth infected host. Trends parasitology 31, 405–406 (2015).

36. Audebert, C. et al. Colonization with the enteric protozoa blastocystis is associated with increased diversity of human gut bacterial microbiota. Sci. reports 6, 25255 (2016).

37. Scanlan, P. D. & Stensvold, C. R. Blastocystis: getting to grips with our guileful guest. Trends parasitology 29, 523–529 (2013).

38. Scanlan, P. D. et al. The microbial eukaryote blastocystis is a prevalent and diverse member of the healthy human gut microbiota. FEMS microbiology ecology 90, 326–330 (2014).

39. Faith, J. J. et al. The long-term stability of the human gut microbiota. Science 341, 1237439 (2013).

40. Coyte, K. Z., Schluter, J. & Foster, K. R. The ecology of the microbiome: networks, competition, and stability. Science 350, 663–666 (2015).

41. Hutzil, S., Sandoval-Motta, S., Frank, A. & Aldana, M. Modeling the role of the microbiome in evolution. Front. physiology 9, 1836 (2018).

42. Roli, A., Villani, M., Filisetti, A. & Serra, R. Dynamical criticality: overview and open questions. J. Syst. Sci. Complex. 31, 647–663 (2018).

43. Goldberger, A. L. Fractal mechanisms in the electrophysiology of the heart. IEEE Eng. Medicine Biol. Mag. 11, 47–52 (1992).

44. Kiyono, K., Struzik, Z. R., Aoyagi, N., Togo, F. & Yamamoto, Y. Phase transition in a healthy human heart rate. Phys. review letters 95, 058101 (2005).

45. Massobrio, P., de Arcangelis, L., Pasquale, V., Jensen, H. J. & Plenz, D. Criticality as a signature of healthy neural systems. Front. systems neuroscience 9, 22 (2015).

46. Rivera, A. L. et al. Looking for biomarkers in physiological time series. In Quantitative Models for Microscopic to Macroscopic Biological Macromolecules and Tissues, 111–131 (Springer, 2018).

47. Goldberger, A. L. et al. Fractal dynamics in physiology: alterations with disease and aging. Proc. national academy sciences 99, 2466–2472 (2002).

48. Derrida, B. & Pomeau, Y. Random networks of automata: a simple annealed approximation. EPL (Europhysics Lett. 1, 45 (1986).

49. Aldana, M. Boolean dynamics of networks with scale-free topology. Phys. D: Nonlinear Phenom. 185, 45–66 (2003).

50. Aldana, M., Balleza, E., Kauffman, S. & Resendiz, O. Robustness and evolvability in genetic regulatory networks. J. theoretical biology 245, 433–448 (2007).

51. Torres-Sosa, C., Huang, S. & Aldana, M. Criticality is an emergent property of genetic networks that exhibit evolvability. PLoS computational biology 8, e1002669 (2012).

52. Langton, C. G. Computation at the edge of chaos: phase transitions and emergent computation. Phys. D: Nonlinear Phenom. 42, 12–37 (1990).

53. Nykter, M. et al. Critical networks exhibit maximal information diversity in structure-dynamics relationships. Phys. review letters 100, 058702 (2008).

54. Kinouchi, O. & Copelli, M. Optimal dynamical range of excitable networks at criticality. Nat. physics 2, 348 (2006).

55. Shmulevich, I., Kauffman, S. A. & Aldana, M. Eukaryotic cells are dynamically ordered or critical but not chaotic. Proc. Natl. Acad. Sci. 102, 13439–13444 (2005).

56. Serra, R., Villani, M., Graudenzi, A. & Kauffman, S. Why a simple model of genetic regulatory networks describes the distribution of avalanches in gene expression data. J. theoretical biology 246, 449–460 (2007).

57. Balleza, E. et al. Critical dynamics in genetic regulatory networks: examples from four kingdoms. PLoS One 3, e2456 (2008).

58. Daniels, B. C. et al. Criticality distinguishes the ensemble of biological regulatory networks. Phys. review letters 121, 138102 (2018).

59. Allesina, S. & Tang, S. Stability criteria for complex ecosystems. Nature 483, 205 (2012).

60. Gershenson, C. & Fernández, N. Complexity and information: Measuring emergence, self-organization, and homeostasis at multiple scales. Complexity 18, 29–44 (2012).

61. Bak, P. & Paczuski, M. Complexity, contingency, and criticality. Proc. Natl. Acad. Sci. 92, 6689–6696 (1995).

62. Christensen, K. & Moloney, N. R. Complexity and criticality, vol. 1 (World Scientific Publishing Company, 2005).

63. Chin, C.-H. et al. cytohubba: identifying hub objects and sub-networks from complex interactome. BMC systems biology 8, S11 (2014).

64. Santamaría-Bonfil, G., Gershenson, C. & Fernández, N. A package for measuring emergence, self-organization, and complexity based on shannon entropy. Front. Robotics AI 4, 10 (2017).

65. Ramírez-Carrillo, E. et al. Assessing sustainability in north america’s ecosystems using criticality and information theory. PloS one 13, e0200382 (2018).

66. Taleb, N. N. (anti) fragility and convex responses in medicine. In International Conference on Complex Systems, 299–325 (Springer, 2018).

67. Campbell, S. J. et al. Complexities and perplexities: a critical appraisal of the evidence for soil-transmitted helminth infection-related morbidity. PLoS neglected tropical diseases 10, e0004566 (2016).

68. DGIS.

69. INEGI. Datos.

70. Caporaso, J. G. et al. Qiime allows analysis of high-throughput community sequencing data. Nat. methods 7, 335 (2010).

71. Carrillo-Araujo, M. et al. Phyllostomid bat microbiome composition is associated to host phylogeny and feeding strategies. Front. microbiology 6, 447 (2015).

72. Allali, I. et al. A comparison of sequencing platforms and bioinformatics pipelines for compositional analysis of the gut microbiome. BMC microbiology 17, 194 (2017).

73. Paulson, J. N., Pop, M. & Bravo, H. C. metagenomeseq: Statistical analysis for sparse high-throughput sequencing. Bioconductor package 1, 63 (2013).

74. Cringoli, G., Rinaldi, L., Maurelli, M. P. & Utzinger, J. Flotac: new multivalent techniques for qualitative and quantitative copromicroscopic diagnosis of parasites in animals and humans. Nat. protocols 5, 503 (2010).

75. Becker, S. L. et al. Toward the 2020 goal of soil-transmitted helminthiasis control and elimination. PLoS neglected tropical diseases 12, e0006606 (2018).

76. Team, R. C. et al. R: A language and environment for statistical computing. (2013).

77. Shannon, P. et al. Cytoscape: a software environment for integrated models of biomolecular interaction networks. Genome research 13, 2498–2504 (2003).

78. Malek, M., Ibragimov, R., Albrecht, M. & Baumbach, J. Cytogedevo—global alignment of biological networks with cytoscape. Bioinformatics 32, 1259–1261 (2015).

79. Ibragimov, R., Malek, M., Guo, J. & Baumbach, J. Gedevo: an evolutionary graph edit distance algorithm for biological network alignment. In German Conference on Bioinformatics 2013 (Schloss Dagstuhl-Leibniz-Zentrum fuer Informatik, 2013).

